# Therapeutic depletion of CD8+ T-cells prevents myelin pathology in Globoid Cell Leukodystrophy

**DOI:** 10.1101/2022.09.30.510367

**Authors:** Pearl A. Sutter, Antoine Ménoret, Evan R. Jellison, Alexandra M. Nicaise, Allison M. Bradbury, Anthony T. Vella, Ernesto R. Bongarzone, Stephen J. Crocker

## Abstract

Globoid cell leukodystrophy (GLD) or Krabbe’s disease is a fatal genetic demyelinating disease of the central nervous system caused by loss-of-function mutations in the galactosylceramidase (galc) gene. While the metabolic basis for disease is known, the understanding of how this results in neuropathology is not well understood. Herein we report that the rapid and protracted elevation of CD8+ cytotoxic T lymphocytes occurs coincident with clinical disease in a mouse model of GLD. Administration of a function blocking antibody against CD8α effectively prevented disease onset, reduced morbidity and mortality and prevented CNS demyelination in mice. These data indicate that subsequent to the genetic cause of disease, neuropathology is driven by pathogenic CD8+ T cells, thus offering novel therapeutic potential for treatment of GLD.

**One-Sentence Summary:** CD8 T-cells mediate demyelination and neuroinflammation in a genetic white matter disease.

---

Globoid Cell Leukodystrophy (GLD) is a fatal demyelinating disease of the central nervous system (CNS) that leads to paralysis and death in 99% of affected children before the age of 5 (*1, 2*). Loss of function mutation in the enzyme galactocerebrosidase (GALC) in GLD leads to an accumulation of galactosylsphingosine (“psychosine”) which is known to contribute to the hallmarks of GLD neuropathology including neuroinflammation and demyelination (*3, 4*). Research has shown that neuroinflammation in GLD precedes demyelination (*5–7*), and we have confirmed evidence of CD8+ T-cells in human GLD brain, as well as identified comparable findings in naturally occurred GLD models in both dogs (fig S1A-S1D), and the twitcher mouse model (*8–11*). However, a functional role for CD8+ T cells had not been identified in this disease.

To better understand the T cell populations within the CNS of *twi* mice we conducted flow cytometry at four timepoints spanning the short 45-day lifespan of *twi* mice which coincided with established phases of disease progression: pre-clinical disease (postnatal (p)14), disease onset (p21), fully developed clinical disease (p30), and end-stage disease (p40) (Fig. 1A, fig S2). A significant increase in CD8+ T cells at the time of disease onset (p21) was identified and found to continue to increase as disease progressed (Fig. 1B–1E). Activated CD4+ T cells were also identified in *twi* CNS at p40, yet their relative numbers were lower than CD8+ T cells and CNS infiltration was not observed until the end stage of disease ~p40 (fig S3A-S3B). Measurement of effector memory CD8+ T-cells (CD8+/CD44+/CD62L-) confirmed that the CNS infiltrating CD8+ T-cells in *twi* mice were highly activated during *twi* disease progression (Fig. 1F–1H), as were the infiltrating CD4+ T-cells at p40 (fig S3C-S3E). In contrast, comparison of each CD8+ and CD4+ effector memory (CD44+/CD62L-) populations from spleens of *twi* and wildtype (wt) littermate controls showed similar levels of activation, which suggested negligible differences in CD8+ T cell activation outside the CNS (Fig. 1I, fig S4A-S4I). Immunohistochemistry on *twi* CNS tissues identified CD8+ T cells in cortex (Fig. 1J) and white matter (myelin basic protein+) (Fig. 1K). A significantly higher number of CD8+ T-cells were found in *twi* compared to wt, with a majority of the CD8+ T-cells being found in the cortex (Fig 1L–1M). No CD8+ T-cells were observed in wt littermates (Fig. 1L).

**Fig. 1.**
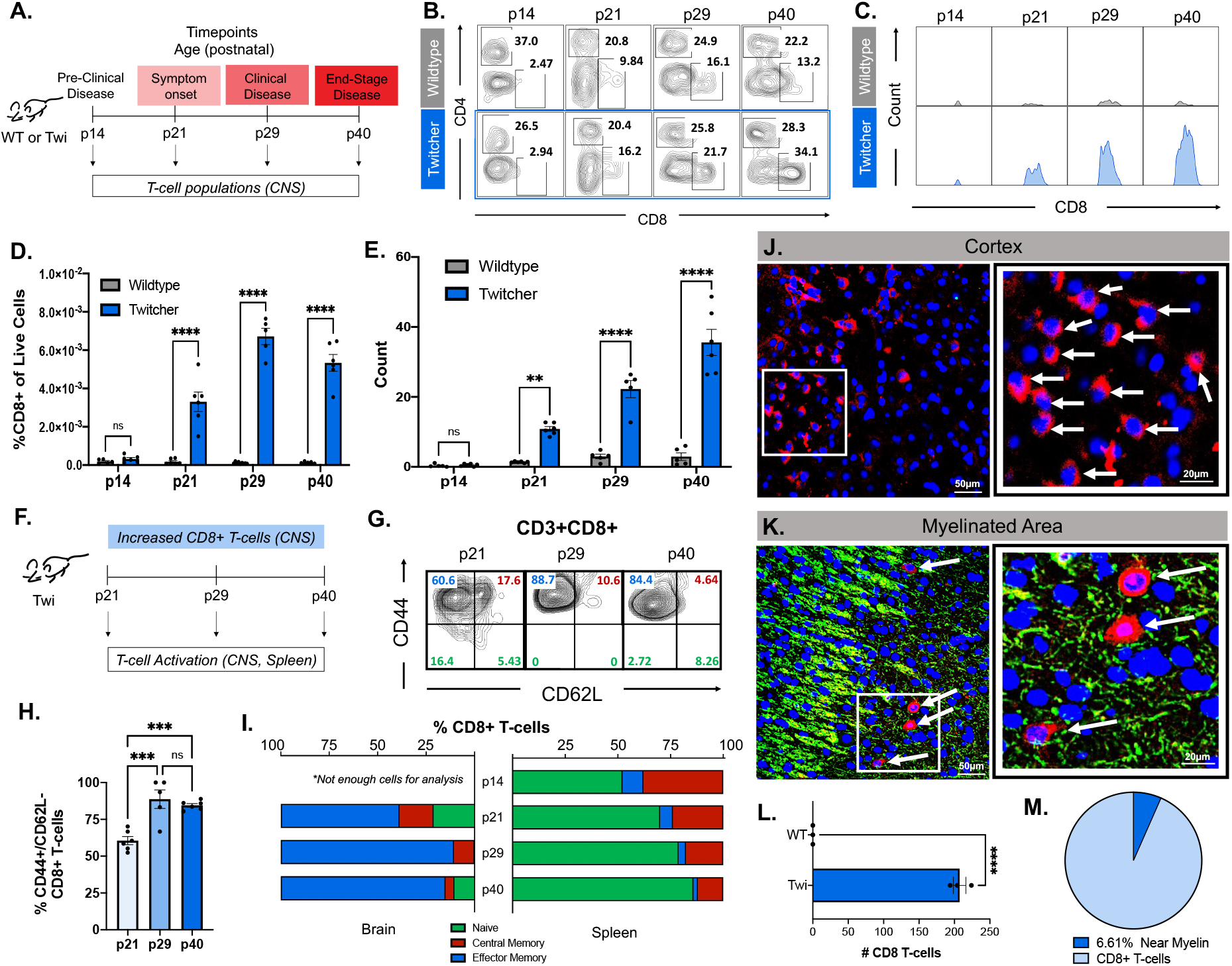
Robust activated CD8+ T-cell populations in the *twi* CNS correlate with disease progression. (**A**) Schematic showing *twi* disease progression with correlated murine postnatal age. (**B**) Flow cytometry plots showing the percentage of CD4+ and CD8+ T-cells (gated on Scatter:Live:CD45^high^:CD3+; fig S1) in WT and *twi* mouse brains at four timepoints during disease progression; n=5-6. (**C**) Histograms showing counts of CD8+ T-cells in WT and *twi* mice during disease progression; n=5-6. (**D**) Percent of CD3+:CD8+ T-cells out of all live cells in WT and Twi CNS; n=5-6. (**E**) Counts of CD8+ T-cells in WT and *twi* mice brains, n=5-6. (**F**) Schematic showing analysis of CD8+ T-cell activation was performed only in *twi* CNS and spleens. (**G**) Representative flow plots of gated *twi* CD8+ T-cells in *twi* CNS gated by CD44 and CD62L to characterize T-cell activation in CNS. (**H**) The percent of effector memory CD8+ T-cells in *twi* CNS increased with disease progression. (**I**) *Twi* CD8+ T-cells in CNS vs spleen show differences in percentages of activated T-cells. Representative image of clustered CD8+ T-cells in *twi* cortex (**J**) and myelinated areas (**K**) at p29. White box indicates area for zoomed in image on right; white arrows point to CD8+ T-cells; red = CD8, green = MBP, blue = DAPI. (**L**) Number of CD8+ T-cells is increased in *twi* CNS at p29 with (**M**) 6.61% of CD8+ T-cells found in myelinated areas, n=10 images/animal (5 cortex, 5 myelinated areas), 3 animals/group. **p<0.01, ***p<0.001, ****p<0.0001; statistical tests used include t-tests (L), 1-way ANOVA (H), 2-way ANOVA (D,E).

To determine the functional importance of CD8+ T-cells to GLD-like disease in the *twi* mouse, CD8+ T cells were depleted by administration of anti-mouse CD8α antibody (*twi*:CD8α; 300μg, i.p) (*12*) every 5^th^ day starting at p14 (Fig. 2A–2C). IgG2b isotype matched antibody (*twi*:Iso-IgG2; 300μg, i.p.) administration did not affect CD8+ T cell counts, nor did CD8α antibody treatment affect CD4+ T-cell populations in *twi* mice (Fig. 2C, fig S5A-S5C). Depletion of CD8+ T-cells in *twi* mice significantly improved overall wellness (Fig. 2D–2E), delayed onset and overall severity of disease (Fig 2D, 2F). Reduced disease severity was evident as 10/10 *twi*: Iso-IgG2 mice developed paralysis yet among *twi*:CD8α mice only 7/9 developed a mild twitching phenotype by p29 as measured by the disease severity scoring (DSS) (*13*) (Fig. 2E–2F). Moreover, when compared with the natural *twi* lifespan of P45, which was used as the experimental endpoint of this study, *twi*:CD8α mice exhibited increased survival (Fig. 2G).

**Fig. 2.**
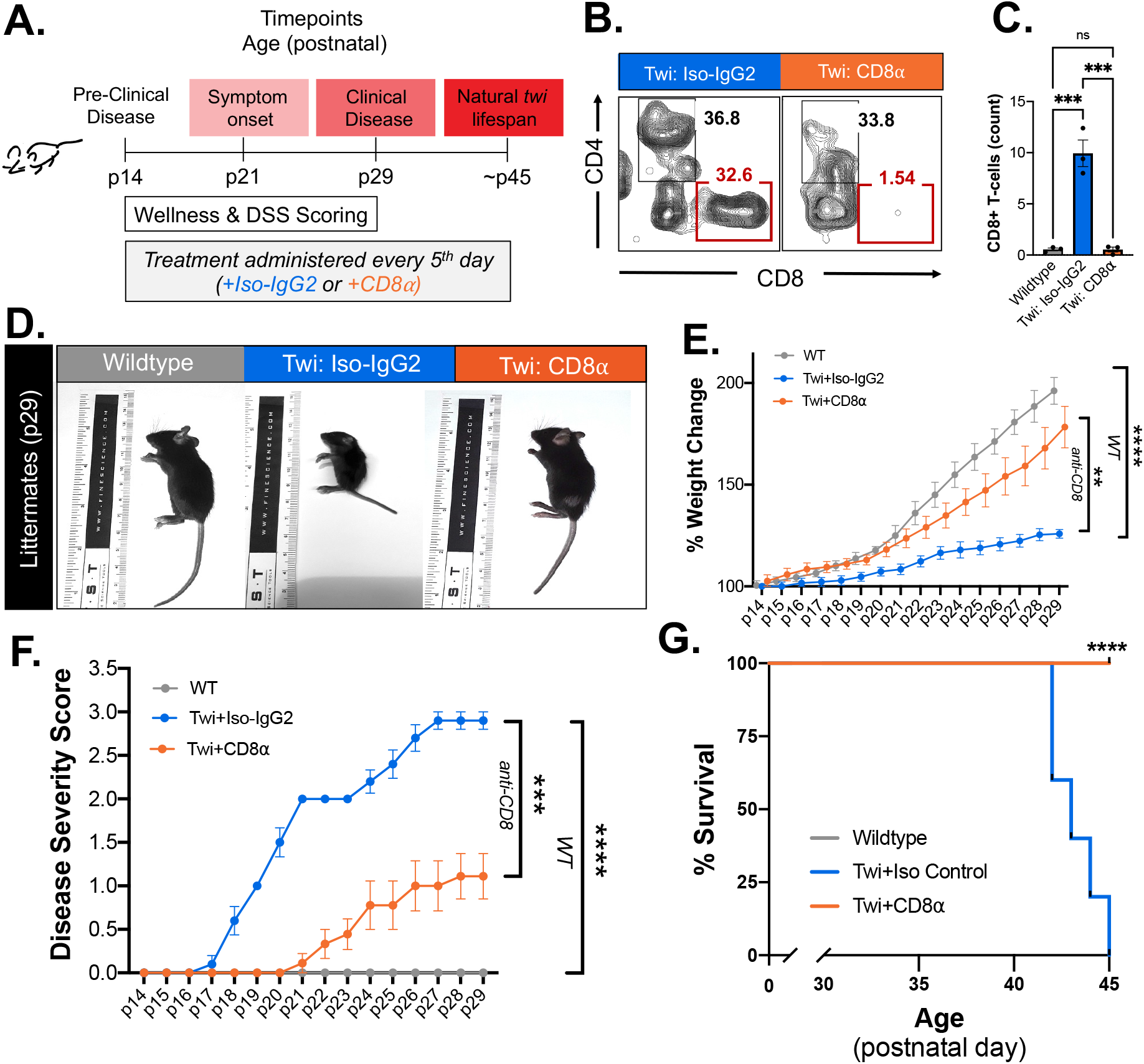
Depletion of CD8+ T-cells decreases clinical disease in twi mice. (**A**) Experimental schematic showing the timing of the CD8α or Isotype IgG control injections in WT and *twi* mice. (**B**) Flow plots of brain CD3+ T-cells (as gated in fig S1) in WT, *twi:Iso-IgG2, twi:CD8a* at p29 show depletion of CD8+ T-cells in *twi:CD8a*., **(C**) Administration of CD8α antibody shows specific depletion of CD8+ T-cells without impacting CD4+ T-cells; n=3. (**D**) Representative mice from each treatment group (WT, *twi:Iso-IgG2, twi:CD8a*) at p29. (**E**) Percent weight change normalized to p14 showing the weight gain of each treatment group between p14 – p29; n=8-10. (**F**) Disease severity scoring (DSS) for each treatment group between p14 – p29; n=8-10. (**G**) Treatment with CD8α extended lifespan of *twi* mice past the normal *twi* lifespan; n=5. **p<0.01, ***p<0.001, ****p<0.0001; statistical tests used include t-tests (C), 2-way ANOVA (E-G).

The impact of CD8+ T cell depletion on CNS demyelination and neuroinflammation was also analyzed (Fig. 3A). Measurement of myelinated axon g-ratios in the corpus callosum of wildtype, *twi*:Iso-IgG2, and *twi*:CD8α mice determined that depletion of CD8+ T cells ameliorated demyelination (Fig. 3B-D) and profoundly decreased the percentage of demyelinated axons from 24.6% in *twi*:Iso-IgG2 to 0.9% in *twi*:CD8α mice (Fig. 3E). These findings were consistent with the preservation of myelin throughout the CNS in *twi*:CD8α mice, as indicated by histological staining of the cerebellum with luxol fast blue (Figure 3F). Neuroinflammation was evaluated by immunohistochemistry staining to examine activated astrocytes (GFAP) and microglia/macrophages (Iba-1). Depletion of CD8+ T-cells in *twi* mice showed reduced activated astrocytes and microglia/macrophages in the cerebellum (Fig. 3G-I), cortex (Fig 3J-L), hippocampus and corpus callosum (fig S6A-S6D), as compared *twi*:Iso-IgG2. These findings were corroborated with RT-qPCR on hemibrains that showed astrocytes (*GFAP*) and microglia/macrophages (*IBA1, CD86*) exhibited reduced activation in *twi*:CD8α mice compared to *twi*: Iso-IgG2, although these markers still remained elevated when compared to wt littermates (Fig. 3M-O). Multiplex cytokine arrays also identified significantly decreased levels of IFNg, IL-2, TNFα, (Fig. 3P-R), and IL-6, IL-1α, CXCL1, CCL2, CCL3, CXCL9, and CCL5 in *twi*:CD8α mice compared with *twi*:Iso-IgG2 mice(fig S7A).

**Fig. 3.**
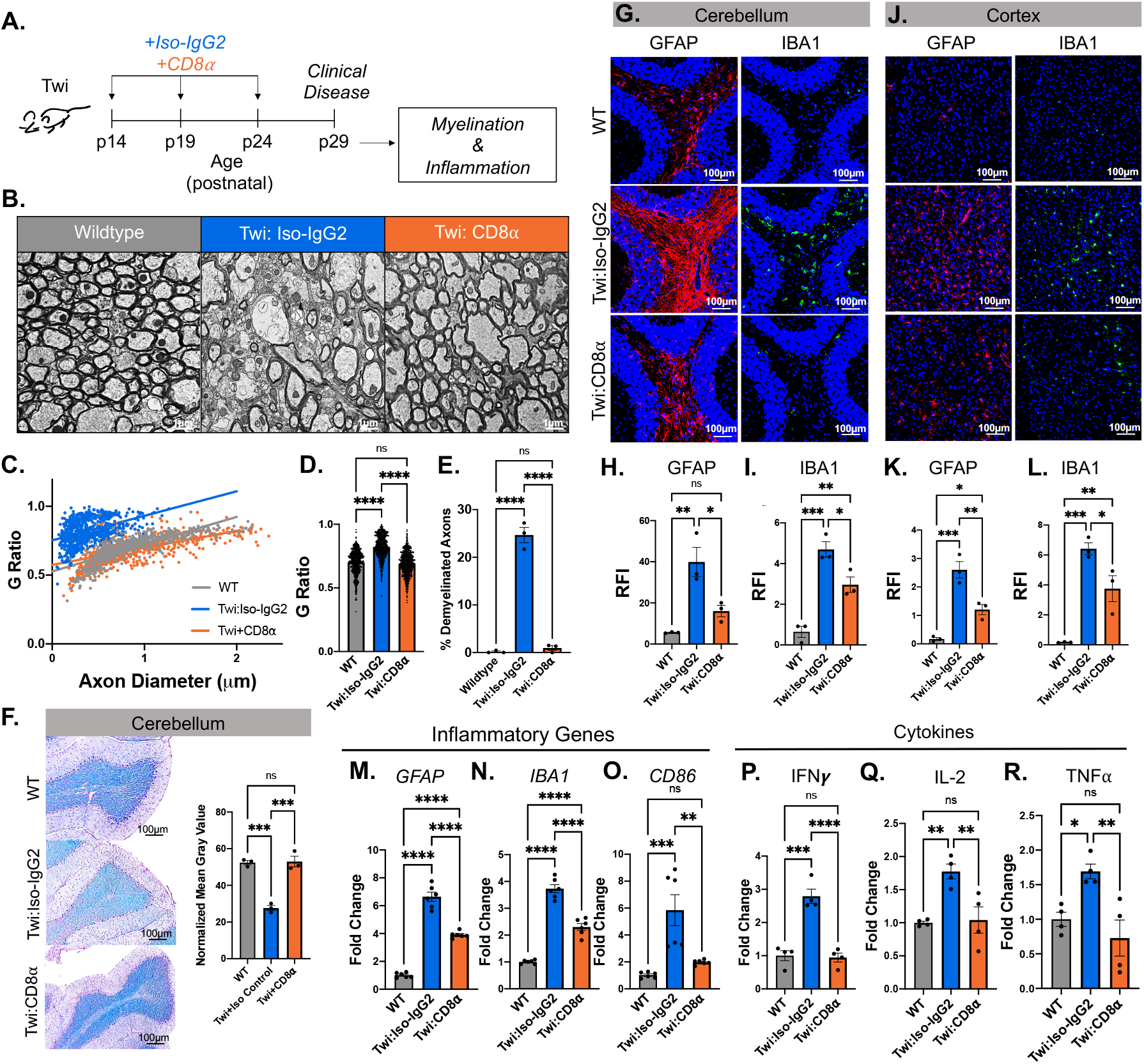
Depletion of CD8+ T-cells ameliorates myelin pathology and attenuates CNS inflammation in twi mice at p29. (**A**) Experimental schematic showing treatment regime and analysis timeline. (**B**) Representative images of transmission electron microscopy (TEM) images of the corpus callosum at p29. (**C**) Plotted G-ratio vs. Axon diameter at p29; n=250 axons/animal, 3 animals/group. (**D**) G-ratios for all treatment groups at p29; n=250 axons/animal, 3 animals/group. (**E**) Percentage of demyelinated axons (g-ratio >0.95) present in each treatment group at p29, n=3. (**F**) Representative images of Luxol Fast Blue staining of the cerebellum at p29 with calculated normalized mean gray value for image; n=5 images/animal (averaged), n=3 animals/group. (**G-I**) GFAP and IBA1 staining in the cerebellum of WT, *twi:IsoIgG2*, and *twi:CD8α* with quantification; n=5 images/animal, 3 animals/group. (**J-L**) GFAP and IBA1 staining in the cortex of WT, *twi:IsoIgG2*, and *twi:CD8α* with quantification; n=5 images/animal, 3 animals/group. (**M-P**) Normalized fold change of gene expression of inflammatory markers known to be upregulated in *twi* mice are reduced with CD8+ T-cell depletion, n=6. (**Q-S**) Normalized fold change (to WT) of cytokines from hemibrains of animals from each treatment group; n=4. *p<0.05, **p<0.01, ***p<0.001, ****p<0.0001; statistical tests used include t-tests (D-R).

To provide a profile of the immunological landscape in the *twi* mouse brain, we performed single-cell RNA sequencing of CNS infiltrating CD45+ cells at p21 (*twi* mice compared to wt littermates; Fig. 4A, 4B). This analysis identified five populations of T-cells including CD4+, CD8+, and ***γ*δ** T-cells (Fig 4B, fig S8A). One CD8+ T-cell population showed a 9-fold increase in CD8+ T cells in *twi* mouse brain compared to wt (Fig. 4C, 4D, fig S8B), further validating our earlier findings and consistent with the reported presence of elevated T cells in *twi* and human GLD CNS (*8–11, 14*). Transcriptomic analysis of this profoundly increased CD8+ T cell population confirmed an activated cytotoxic T-cell phenotype indicated by increased expression of cytotoxic markers including *CD160, XCL1, GZMB*, and *CXCR6* (Fig. 4E). *Twi* CD8+ T-cells also had increased expression of many effector molecules and cytokines as well genes involved in the IFN***γ*** response (Fig. 4F) (*15*). Further analysis revealed that CD8+ T-cells in *twi* brain differentially expressed 81 genes, of which 67 were significantly up-regulated (Fig. 4G). Gene ontology analyses of the upregulated genes revealed *twi* CD8+ T-cell involvement in biological and reactome pathways including *T cell Receptor (TCR) signaling, T cell activation, T cell differentiation, positive regulation of leukocyte mediated cytotoxicity*, and several pathways involving antigen presentation such as *antigen processing and presentation via MHC class Ib* and *co-stimulation by the CD28 family* (Fig. 4H). These data provide a basis for evaluating potential underlying mechanism of CD8+ T-cell activity in *twi* mice.

**Fig. 4.**
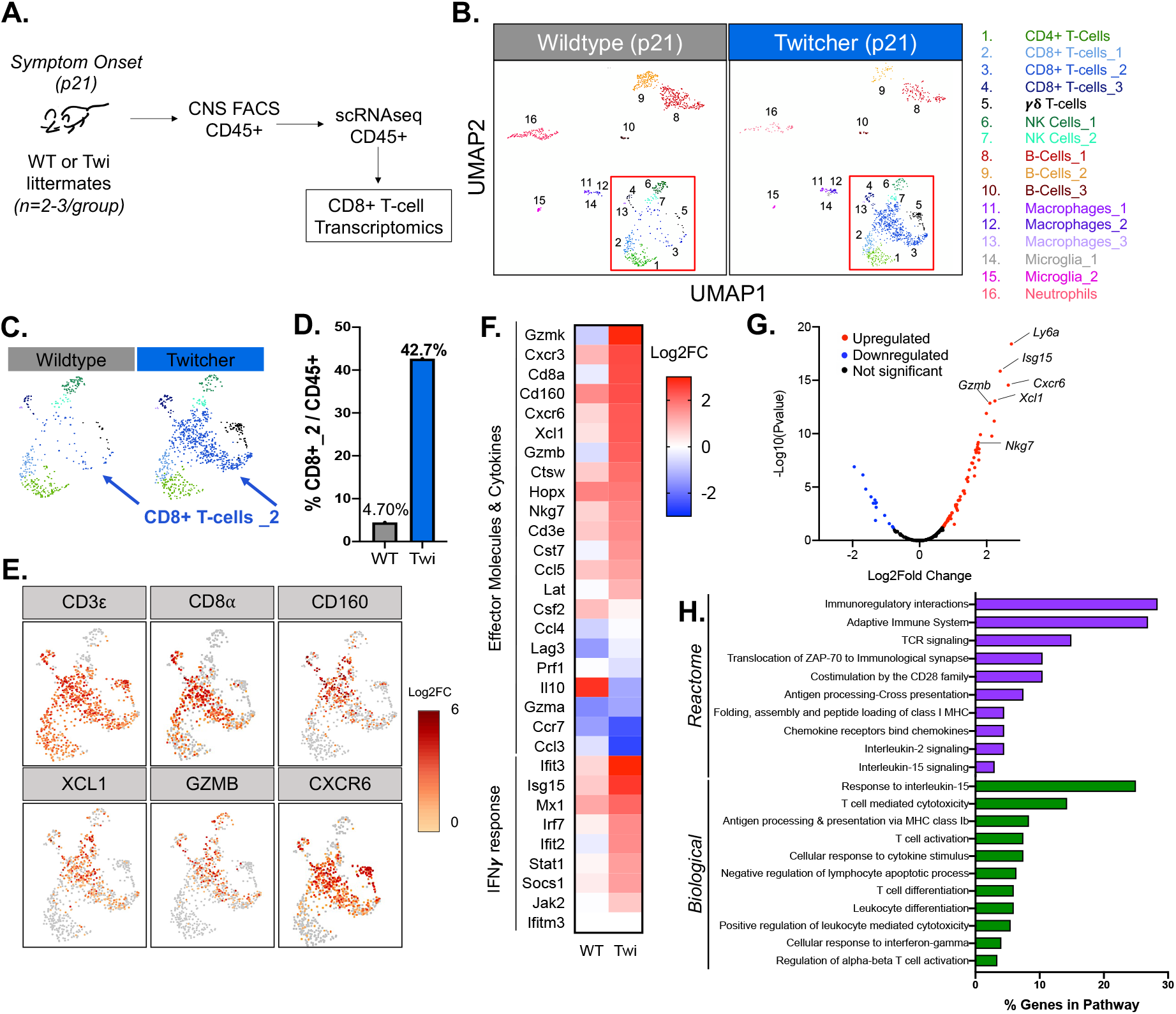
Transcriptomic analysis of CD8+ T-cells in twi CNS at disease onset (p21). **(A)** Experimental workflow for isolating CNS CD45+ cells for analysis by scRNAseq, n=2-3/group. (**B**) UMAP visualization of scRNAseq data of CD45+ populations from p21 WT and *twi* mice shows 16 unique populations. (**C-D**) The *twi CD8+T-cells_2* population shows a 9-fold increase in size compared to the WT CD8+T-cells_2 population. (**E**) UMAP plots showing cytotoxic genes are highly expressed by *twi* CD8+T-cells. (**F**) Effector molecules, cytokines, and genes involved in the IFN***γ*** response show increased expression in *twi* CD8+ T-cells compared to wt. (**G**) Volcano plot showing differentially expressed genes (DEGs) in *twi* CD8+ T-cells compared to wt (upregulated = red; downregulated = blue). (**H**) Significant GO Ontology analysis results for Biological and Reactome processes analyzing the pathways associated with the 67 upregulated genes.

Herein, we have defined a novel role for CD8+ T cells in the initial development of GLD disease and myelin pathology. We have identified that there is spontaneous development of a robust, pathological CD8+ T cell population that is associated with GLD disease in this naturally occurring disease model. It is currently not known how the metabolic consequences of *galc* mutations in GLD result in the development of such a profound immunological response. CD8+ T cells are a predominant cytotoxic lymphocyte population found in human neuroinflammatory diseases affecting central nervous system white matter, including GLD (*11, 16–18*). Accordingly, additional study on the genesis of CD8+ T cell responses in this disease model may contribute to our fundamental understanding of how these immunological responses may begin. It is important to point out that this model has some striking parallels and differences from other known autoimmune disease models. First, CD8+ T cell responses in *twi* mice were spontaneous and occurred very early in postnatal development. This naturally occurring CD8+ T-cell mediated phenotype distinguishes it from most other widely adopted models of CNS demyelination often associated with either Multiple Sclerosis (MS) or Neuromyelitis Optica (NMO) (*19*), such as experimental autoimmune models, wherein inoculation with a known peptide or virus results in measurable antigen-specific responses. In contrast to these other models, the antigen specificity and clonality of CD8+ T cells in GLD are not presently known. Interestingly, the rapid and robust development of CD8+ T cells within the CNS compartment of *twi* mice was not preceded by vigorous changes in peripheral CD8+ T cell populations. This observation may suggest a fundamental difference in where cells become highly activated and may engage antigen presenting cells (APCs), potentially including atypical APCs such as glial cells (*20*), which are themselves robustly activated in GLD.

Our findings also portend potential therapeutic benefits for the treatment of GLD by therapeutics which target T cells. For example, some disease modifying treatments for MS have established applications for treating pediatric patients that could then have application for GLD as well (*21, 22*). While we have demonstrated benefit to GLD disease in *twi* mice by affecting CD8+ T-cells, the long-term impact of this approach has yet to be determined. Nevertheless, the effect of this approach on early GLD neuropathology may represent a means by which to enhance existing therapies. For instance, the current therapy for GLD, hematopoietic stem cell transplantation (HSCT) (*23*), is limited by a very narrow window for treatment, often within the first month of life (*24*). Since newborn screening for GLD is not widely available (*25, 26*), new therapeutic approaches such as immunotherapies may be warranted. Applying immunotherapies could extend the therapeutic window for HSCT (*27*) or enhance long-term outcomes from interventions such as gene therapy (*28, 29*), and therein provide greater benefit to more patients. Overall, our findings increase our understanding of GLD pathology and suggest insights for current and novel therapeutic approaches to GLD. Furthermore, these data support *twi* mice as a new model to understand the etiology of CD8+ T cells which affect the CNS, with potential relevance for other diseases.

## Acknowledgments

We also want to thank Dr. Jaime Imitola and Dr. Robert Clark for critical feedback on the manuscript. We are also grateful to Maya Yankova in the UConn School of Medicine Electron Microscopy Core Facility for her expert assistance with transmission electron microscopic analyses of myelin.

## Funding

This work was supported in part by grants from National Institutes of Health National Institutes of Health (R56NS099359-01A1 to SJC) and the National Multiple Sclerosis Society (RG-1802-30211 to SJC). PAS was supported by an NIH F30 award (NS129238-01).

## Author contributions

Each author’s contribution(s) to the paper should be listed [we encourage you to follow the CRediT model]. Each CRediT role should have its own line, and there should not be any punctuation in the initials.

Conceptualization: SJC, PAS, ERB, ATV

Methodology: PAS, AM, EJ, AMB, AMN

Investigation: PAS, AM, SJC

Visualization: PAS, AM, SJC

Funding acquisition: SJC, PAS, ERB, ATV

Supervision: SJC

Writing – original draft: PAS, SJC

Writing – review & editing: PAS, AM, ERJ, AMN, AMB, ATV, ERB, SJC

## Competing interests

Authors declare that they have no competing interests.

## Data and materials availability

Data available on request from authors.

## Supplementary Materials

Materials and Methods

Figs. S1 - S7

Tables S1 - S2

References (*30–34*)

## Notes

### Competing Interest Statement

The authors have declared no competing interest.

