## Supplemental Materials for "Therapeutic depletion of CD8+ T-cells prevents myelin pathology in Globoid Cell Leukodystrophy"

**This PDF file includes:**

Materials and Methods
Supplementary Text
Figs. S1 to S7
Tables S1 to S2
Captions for Data S1 to S2

**Other Supplementary Materials for this manuscript include the following:**

Data S1 to S2

**Materials and Methods**

Animals: All procedures involving animals were conducted with approval from the Institutional Animal Care and Use Committee at the University of Connecticut School of Medicine in accordance with guidelines set forth by the National Research Council of the National Academies Guide for the Care and Use of Laboratory Animals. Mice used in this study were wildtype C57BL/6J (strain #000663) and twitcher C57BL/6J (strain #000845). Mice used in this study were litter matched pups with confirmed genotypes using the following primers (Forward: 5'-GCTTGGAATTTGGTGGCTAG-3'; Reverse: 5'-GATGGTGAGGTTTCCCAAGC-3') *(30).* Dog tissue used in this study was kindly received from co-author Dr. Allison Bradbury *(31, 32).*

Human Brain Tissues: GLD and age-matched control patient (1-2 years old) paraffin-embedded brain tissue was obtained from the National Institute of Child Health and Human Development (NICHD) Brain and Tissue Bank for Developmental Disorders (Baltimore, MD).

CNS Flow Cytometry: At the desired timepoint, animals were euthanized using isoflurane and then received a cardiac perfusion with 1X PBS. Hemibrains at each timepoint were collected and then enzymatically dissociated using collagenase D (12.5 mg/ml) and DNase I (1mg/ml) for 30 minutes at 37°C. CNS tissue was resuspended into 10 ml HBSS+ containing 10% fetal bovine serum (FBS) to neutralize enzymatic activity. Tissue was then strained with a 70µm cell strainer and centrifuged at 1000 RPM for 5 minutes at 4°C. The pellet was resuspended in 30% Percoll, carefully underlaid with 70% Percoll, then spun at 300 x g for 23 minutes at 4°C. Myelin debris was removed from the top layer and leukocytes were collected from the interface between the 30% and 70% Percoll layers. Collected leukocytes were resuspended in 10% FBS in HBSS+, centrifuged at 1000 RPM for 5 minutes at 4°C, and then resuspended in the appropriate antibody cocktail for 20 minutes at 4°C. For samples prepared for single-cell RNA sequencing (scRNAseq), each sample was stained with primary conjugated CD45 antibody and sorted into CD45+ and CD45- cell populations. All other samples for flow cytometry stained with primary conjugated antibodies for CD45, CD3, CD4, CD8a, CD62L, and CD44 (Table S1) for immunophenotyping by flow cytometry. Samples were then resuspended in 1 mL of 1X PBS, centrifuged, and resuspended in the 200µL of 1X PBS for flow analysis on LSR II (Becton Dickinson). Single color compensation controls and FMO controls were prepared for each experiment to account for differences in detector sensitivity across experimental timepoints. Figure S1 shows the gating strategy for identification of T-cell populations.

Spleen Flow Cytometry: At the desired timepoint, animals were euthanized using isoflurane and then received a cardiac perfusion with 1X PBS. Spleens were removed and crushed through a 70µm cell strainer, washed with 1X PBS, then centrifuged at 1000 RPM for 4 minutes. Samples were resuspended in 5 mL of Red Blood Cell Lysis Buffer for 5 minutes at RT, washed with 45 mL of 1X PBS, and centrifuged at 1000 RPM for 4 minutes. Cells were counted and $1x10^5$ cells were isolated from each spleen for staining. Spleen samples were stained and analyzed as described above in the CNS FACS methods section.

Histology and Immunohistochemistry: At p29 mice were perfused with 10mL of 4% paraformaldehyde and then hemibrains were left to fix in 4% paraformaldehyde for 24 hours. Tissues were embedded in paraffin and then sliced into 8µM sections and placed on superfrost plus glass slides. All sections (mouse, dog, and human) were heated for 30 minutes at 60°C, then rehydrated using 3 washes of 100% xylene for 5 minutes each, followed by 5 minute washes in a decreasing ETOH gradient (100%, 95%, 70%). Slides were then washed with 1XPBS and $diH_2o$ and stained either with LFB or prepared for prepared for immunohistochemistry (IHC) by performing antigen unmasking in 0.01M citrate acid buffer. Slides for IHC were blocked for 1 hour at RT in blocking buffer (0.1% Tween, 10% NGS, in 1X PBS), then incubated in primary antibodies (in blocking buffer) overnight at 4°C (see Table S1 for antibody details). Following overnight incubation, slides were washed 5 times for 5 minutes each with 1X PBS and incubated for 1 hour at RT with the appropriate secondary antibody (in blocking buffer). Slides were washed 5 times with 1X PBS, counterstained with DAPI for 5 minutes, then washed and mounted. Sections stained with LFB were imaged at 10x with Olympus IX71 and processed using ImageJ software (National Institutes of Health). Each image was converted to grayscale, then the mean gray value was measured and normalized to the background of each image. IHC labeled sections were imaged with Leica Thunder Microscope and relative fluorescence intensity (RFI) was processed using Image J software (National Institutes of Health).

<u>In vivo Injections and Disease severity Scoring:</u> Twitcher mice received either anti-CD8α antibody (Twi:CD8α; 300µg, i.p.) or isotype matched control antibody (Twi:Iso-IgG2; 300µg, i.p.) every 5$^{th}$ day starting at p14. Mice underwent daily clinical scoring which included weight measurements and evaluation of tremor and locomotion. Weight loss, tremor, and locomotion severity calculated the Disease Severity Score, an established method for measuring Twitcher disease progression (*3*).

<u>Transmission Electron Microscopy</u>: At p29, mice were perfused with 10 mL of fixative containing 2% PFA and 2.5% Glutaraldehyde in 0.1M Cacodylate Buffer (in $diH_2O$). The entire brain was isolated, then incubated in fixative for 1 hour at room temperature (RT). Following postfixing, a 1 mm slice of corpus callosum from each sample was isolated, washed with 0.1M cacodylate buffer and then further fixed in 1% OsO4, 0.8% Ferricyanide in 0.1M cacodylate buffer at RT for 1 hour. Samples were washed 5x with diH20, then underwent *en block staining* in 1% uranyl acetate (in $diH_20$) at RT for 1 hour. Samples were then dehydrated using 10 minute washes of an increasing ETOH gradient (50% ETOH, 75% ETOH, 95% EOTH) at RT, then washed with 3-10 minute washes of 100% ETOH at RT. Samples were next washed with propylene oxide (PO) twice for 5 minutes each, then underwent resin infiltration using PolyBed812 with DMP-30 2, 4, 6-Tris (dimethylaminomethyl) phenol (DMP-30) mixed with differing ratios of PO:Resin (first 1:1 for 1 hour at RT, then 1:3 overnight at RT). Finally, samples were incubated for 24 hours in 100% resin, then embedded and polymerized at 60°C for 48 hours. Each sample was cut into ultra-thin 70 nm sections, placed on grides, and postfixed in PO. Samples were visualized using the Hitachi H7650 transmission electron microscope and all g-ratio and axon diameter measurements were calculated using ImageJ Software (National Institutes of Health).

<u>Quantitative Real-Time Polymerase Chain Reaction (qRT-PCR)</u>: Total RNA was isolated from saline-perfused unfixed hemibrains from p29 mice using TRI Reagent (Sigma) that was added to each sample according to the manufacturer's protocol. cDNA was amplified from isolated RNA via reverse transcription (iScript cDNA synthesis kit, BioRad) and qPCR was performed using specific validated primer pairs for *GFAP, IBA1,* and *CD86* (Table S2) (Integrated DNA Technologies, Coralville, IA) and SsoAdvanced Universal SYBR Green Supermix (BioRad). CFX Connect Real-Time PCR Detection System (BioRad) was used to analyze the amplified cDNA.

Primers for β-actin were used as the housekeeping gene to normalize gene expression among samples. The relative expression of target RNA was calculated using the comparative cycle threshold analysis (ΔΔCT).

CNS Tissue Isolation for Cytokine Array: Mice were perfused with 10mL of ice-cold PBS and whole brain was extracted. Hemibrains were added to glass beads and RIPA Lysis buffer with protease inhibitors, the underwent homogenization for 60 seconds. The homogenate was collected, then centrifuged at 12000 x g for 10 minutes at 4°C to remove insoluble material. The protein concentration of each sample was determined using a Bicinchoninic acid (BCA) assay, then normalized. The cytokines were analyzed using a membrane-based cytokine array (Mouse Inflammation Antibody Array C1; Raybiotech) that analyzes 40 inflammatory cytokines. All material used were from this kit. Briefly, the membranes were blocked with blocking buffer, then 1 mL of each sample was added to each membrane and left to incubate overnight at 4°C. The membranes were then washed 3x each with the wash buffers provided, followed by a 2-hour incubation with biotinylated antibody cocktail at RT. The washes were repeated and then the membranes were incubated with HRP-Streptavidin for 2 hours at RT. Following another wash set, membranes were incubated with detection buffer for 2 minutes, then imaged using chemiluminescence. Data was extracted using ImageJ to obtain spot signal densities from each image, then analyzed using the analysis workbook provided by Raybiotech.

Single Cell RNA Sequencing (scRNAseq): CD45+ flow cytometry sorted (FACS) cells were sent to Jackson Laboratory for scRNAseq processing. Briefly, cells were bar-coded with unique identifiers using lyso-modified oligonucleotides, processed, and visualized using the 10X Genomics Platform. Clusters visualized using UMAP were identified using established cell-type markers.

GO Ontology Analysis: The DEGs *(P<0.05)* identified by scRNAseq between the WT and *twi* CD8+ T-cell populations were collected. We analyzed this gene list using the PANTHER Pathway analysis tool (http://pantherdb.org/) linked through the Gene Ontology Consortium's online database (http://www.geneontology.org/) *(33, 34)*. Gene names were copied into the PANTHER Pathway analysis tool, and organism (*mus musculus),* specific enrichment analysis (*Biological* or *Reactome*), statistical test type and correction (*Fisher's Exact* with *calculation of FDR)* were selected. Significant results were identified by filtering by *false discovery rate (FDR < 0.05)* and *P-value (pval< 0.01)*.

Statistical Analysis: Appropriate statistical analysis were performed for each experiment using GraphPad Prism version 9 for Mac OS X (GraphPad Software). Differences were considered significant when $P < 0.05$. Data are presented as mean +/- SEM.

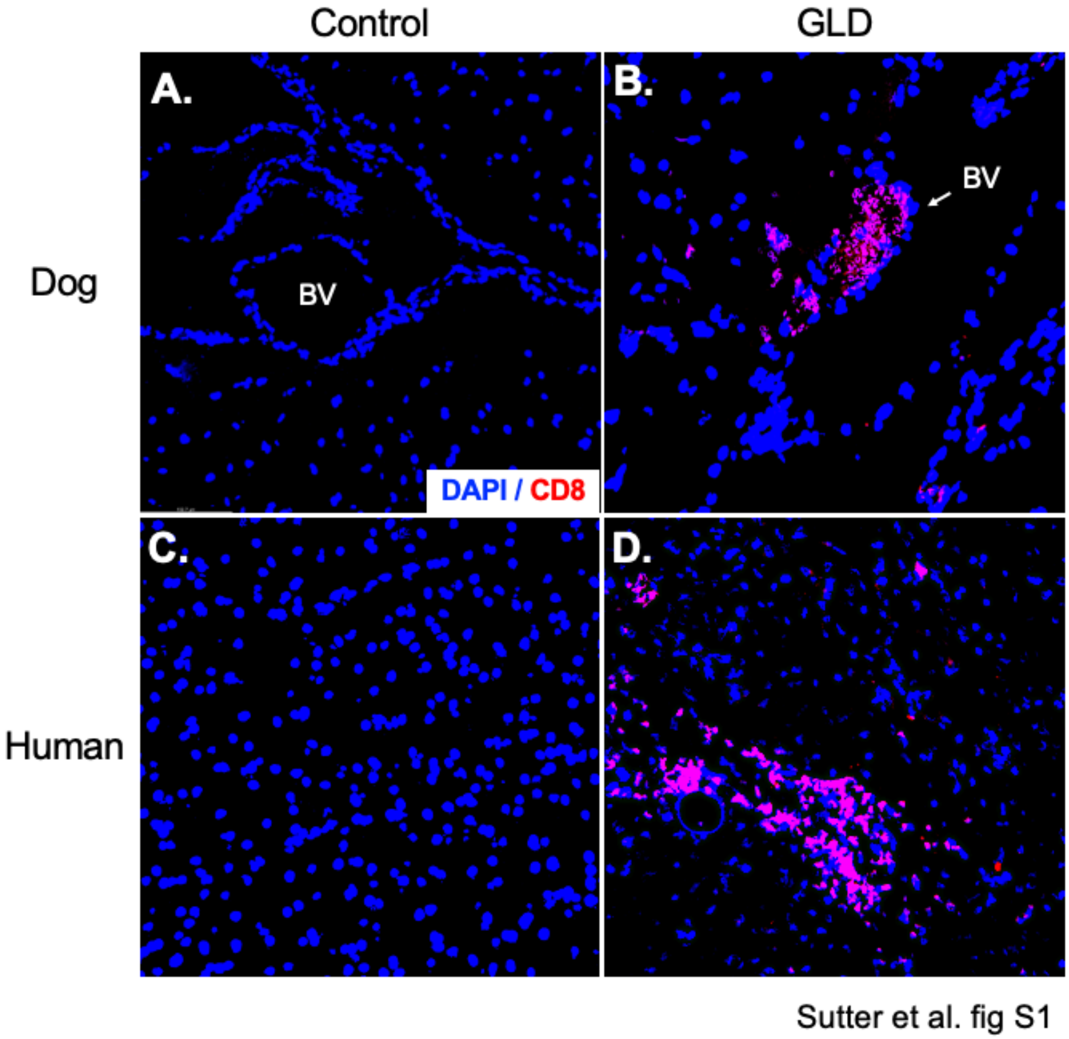


**Fig. S1. CD8+ T-cells are present in the CNS of dog and human GLD.** (**A-B**) CD8+ T-cells infiltrating CNS tissue in dog GLD from blood vessels (BV). (**C-D**) CD8+ T-cells clusters are present in CNS white matter of infantile human GLD.

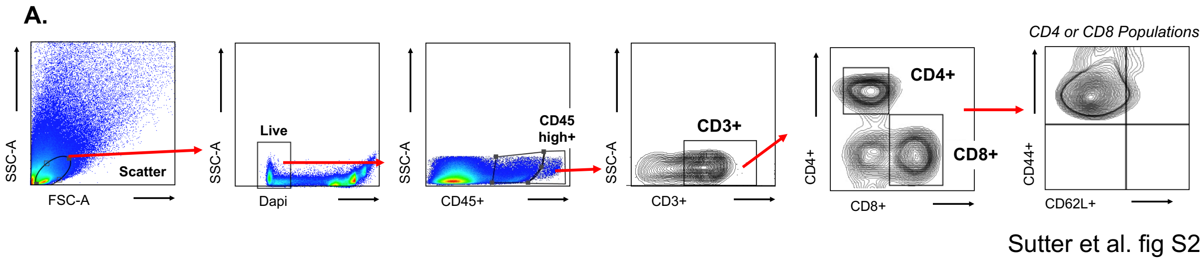


**Fig. S2. Gating Strategy for Leukocyte identification in murine CNS.** (**A**) CNS Flow cytometry gating strategy for identification and analysis of T-cell populations.

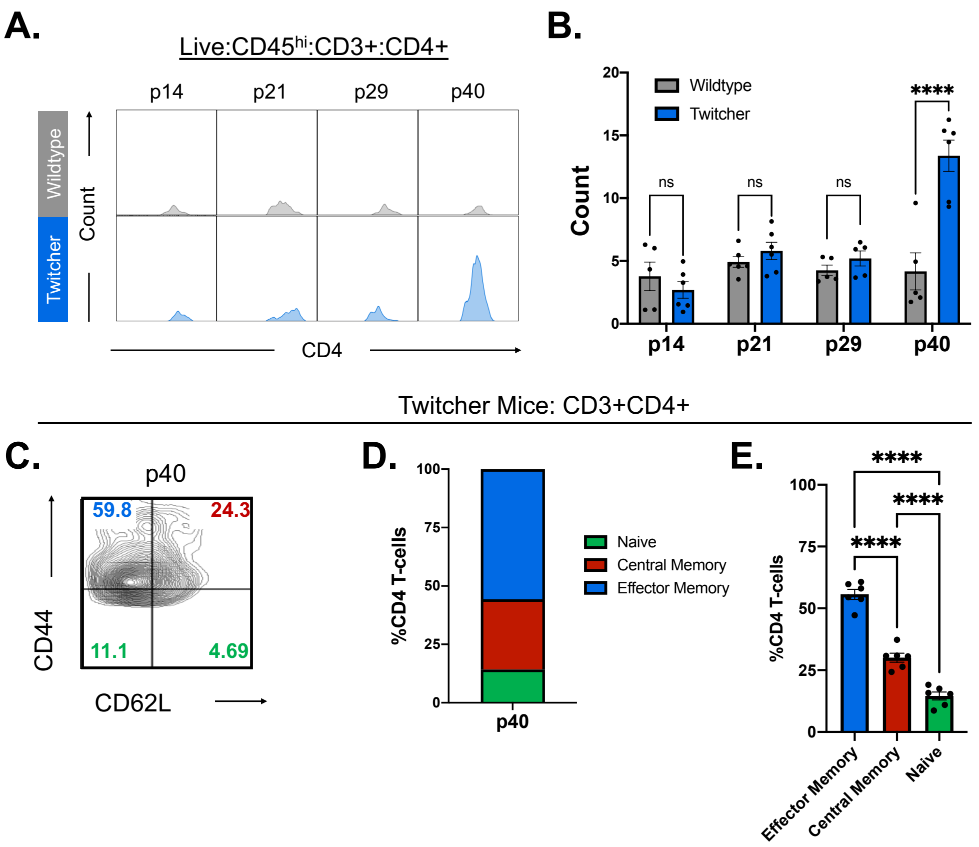


Sutter et al. fig S3

**Fig. S3. Flow cytometry analysis of CD4+ T-cells in *twi* mice CNS during disease progression shows influx at p40.** (**A**) CD4+ T-cells (gated as seen in fig S1) increase in *twi* mice at end stage disease (p40) as shown by representative histograms. (**B**) Area under the curve calculated from the histograms (Fig. S2A) to quantify the CD4+ T-cell population in WT and *twi* mice, n=5-6**.** (**C**) Representative flow plot of gated *twi* CD4+ T-cells gated by CD44 and CD62L to characterize T-cell activation**.** (**D-E**) Analysis of activation states of *twi* CD4+ T-cells shows a majority of the CD4+ T-cells present in *twi* CNS at p40 are activated, n=5-6. ****p<0.0001; statistical tests used include 1-way ANOVA (E), 2-way ANOVA (C)**.**

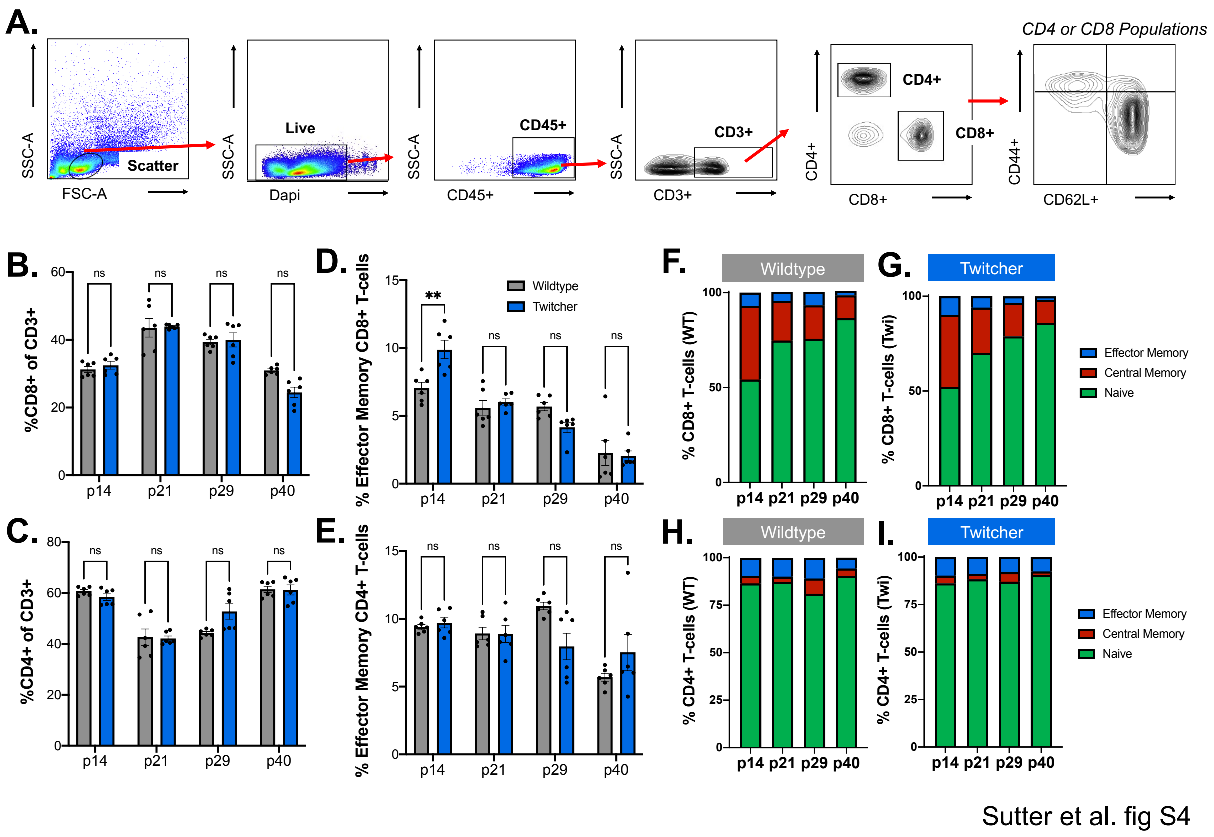


**Fig. S4. Twitcher spleens do not show a robust T-cell response or expansion.** (**A**) Flow cytometry gating strategy for identification of T-cell populations. Percentages of CD8+ cells (**B**) and CD4+ cells (**C**) out of CD3+ T-cells did not differ between WT and *twi* mice during *twi* disease progression, n=6. (**D**) CD8+ effector memory T-cells (CD44+/CD62L-) were elevated at p14 in *twi* mice, but showed no differences at p21, p29, or p40, n=6. (**E**) WT and *twi* CD4+ effector memory T-cell (CD44+/CD62L-) populations did not differ during *twi* disease progression, n=6. Characterization of CD8+ (**F,G**) and CD4+ (**H, I**) T-cell activation in WT and *twi* mice show similar patterns for naïve (CD44-/CD62L+/-), central memory (CD44+/CD62+), and effector memory percentages (CD44+/CD62L-), n=6. **p<0.01; statistical tests used include 2-way ANOVA (B-E).

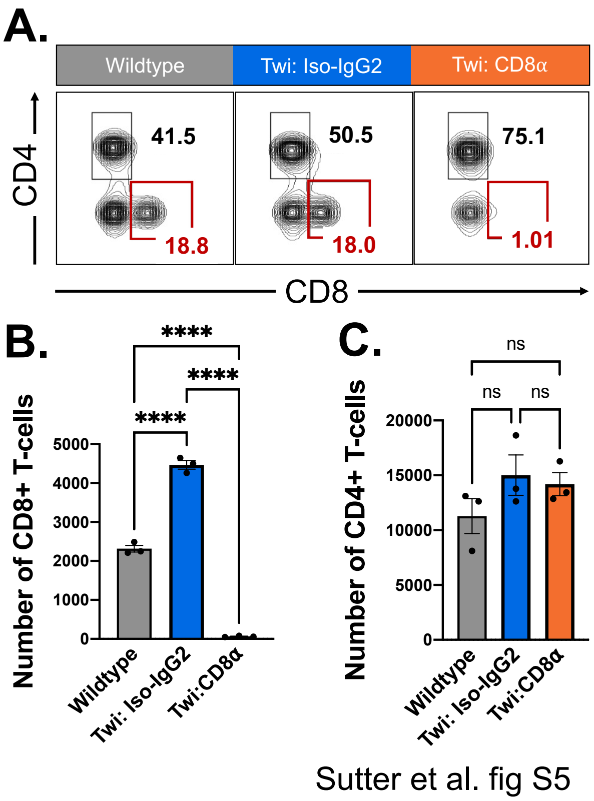


**Fig. S5. Depletion of CD8+ T-cells occurred in the spleen.** (**A**) Representative flow plots of CD3+ T-cells for WT, *twi:Iso-IgG2*, and *twi:CD8α* spleens at p29 show depletion of CD8+ T-cells. (**B**) CD8+ T-cell depletion was specific to CD8+ T-cells (B) without impacting the (**C**) CD4+ T-cell population, n=3. ****p<0.0001; statistical tests used include t-tests (B,C).

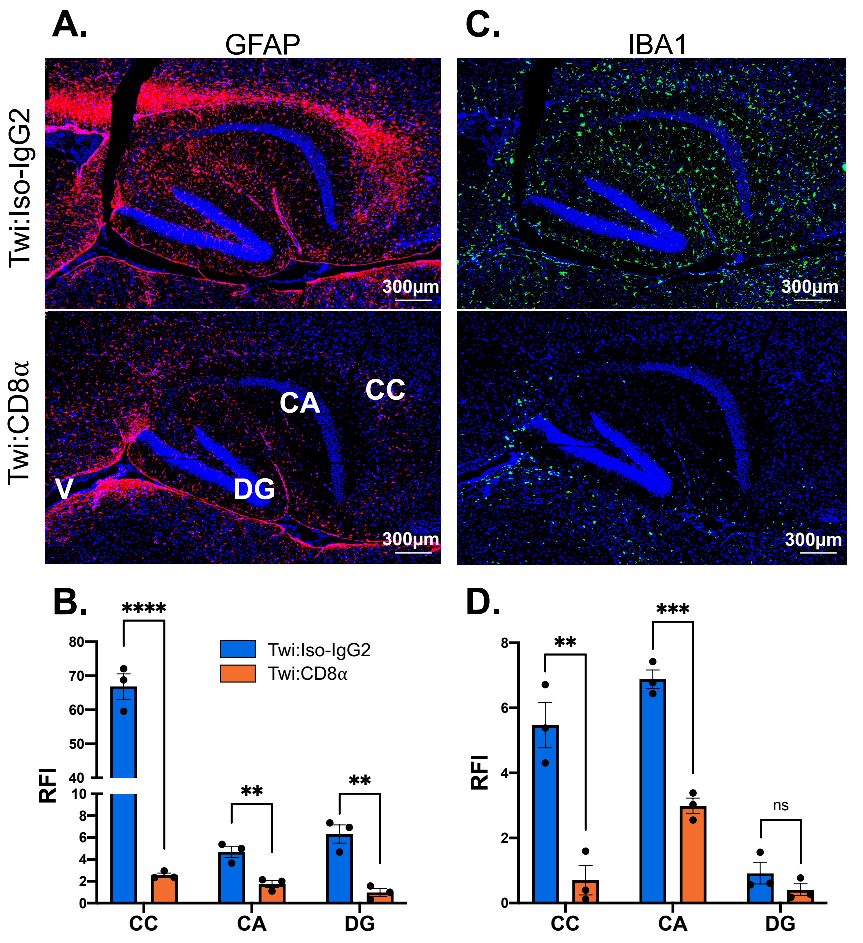


Sutter et al. fig S6

**Fig. S6. Activated astrocytes and macrophages/microglia are reduced in the *twi* hippocampus and corpus callosum following CD8 T-cell depletion.** (**A-B**) Activated astrocytes (GFAP) are decreased in *twi* corpus callosum (CC), cornu ammonis (CA), and dentate gyrus (DG) following CD8+ T-cell depletion, n=5 images/animal averaged, n=3 animals. (**C-D**) Activated microglia/macrophages (IBA1) are decreased in *twi* CC, CA, and DG following CD8+ T-cell depletion, n=5 images/animal averaged, n=3 animals. **p<0.01, ***p<0.001, ****p<0.0001; statistical tests used include t-tests (B,D).

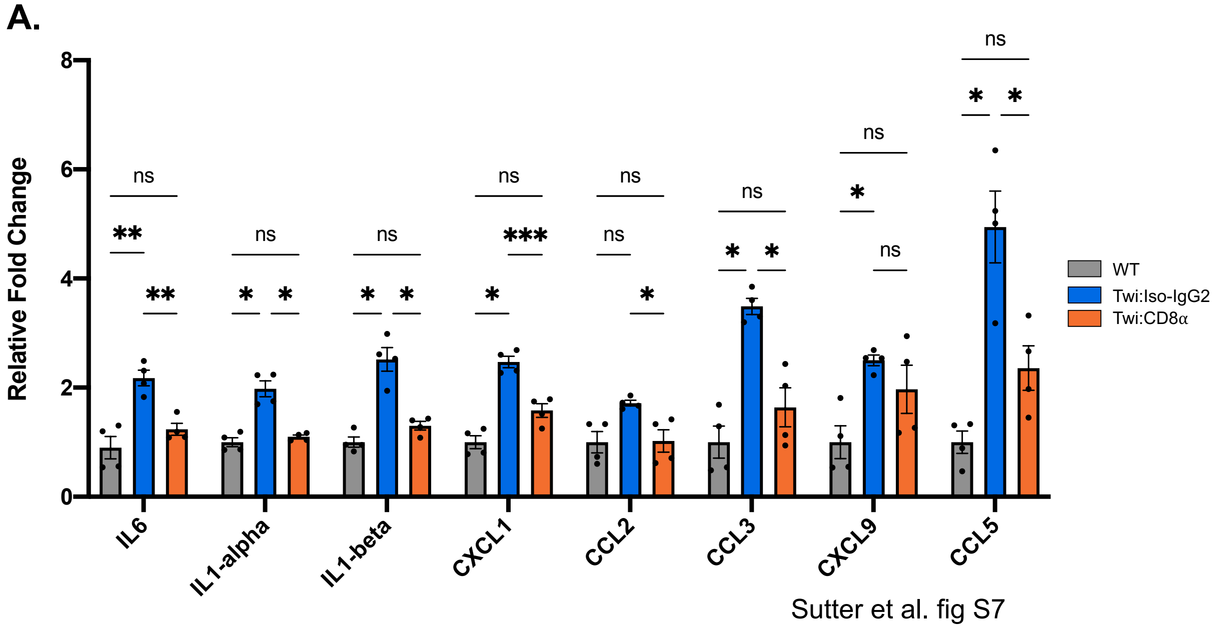


**Fig. S7. CNS cytokines measured from CD8+ depleted animals showed a reduction in inflammatory cytokines known to be elevated in *twi* mice.** (**A**) Fold change of additional cytokines in *twi* CNS show a decrease following CD8+ T-cell depletion in *twi* mice (normalized to WT) n=4. *p<0.05,**p<0.01, ***p<0.001; statistical tests used include 2-way ANOVA (A).

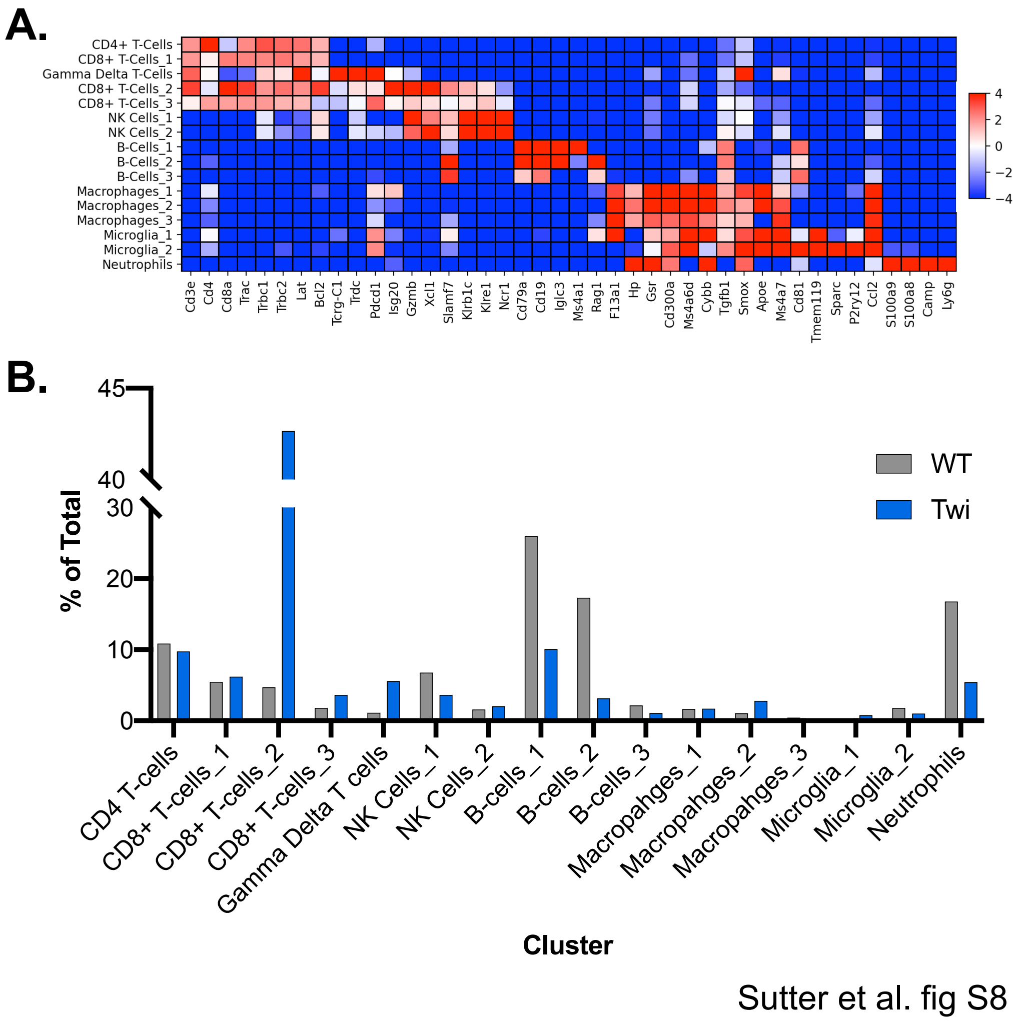


**Fig. S8. scRNAseq clustering and population percentages for WT and Twitcher mice at p21.** (**A**) Heatmap showing gene expression for cluster characterization. (**B**) Percentages of total cells in scRNAseq for each identified cluster in WT and *twi*.

**Table S1. List of Antibodies**

| Antibody | Concentration | Source | Clone |
|---|---|---|---|
| Flow Cytometry | | | |
| CD45 (APC) | 1:200 | BioLegend | 30-F11 |
| CD3 (PE/Cy7) | 1:200 | BioLegend | 145-2C11 |
| CD4 (PE) | 1:200 | BioLegend | GK1.5 |
| CD8a (APC/Cy7)) | 1:400 | BioLegend | 53-6.7 |
| CD44 (V500) | 1:400 | BD Horizon | IM7 |
| CD62L (PE-594) | 1:200 | BD Horizon | MEL-14 |
| DAPI | 1:10,000 | Roche | - |
| Immunohistochemistry | | | |
| MBP | 1:200 | Chemicon | 0603023871 |
| CD8α | 1:500 | BioXCell | 2.43 |
| CD8 | 1:100 | Abcam | - |
| GFA (conjugated to Cy3) | 1:400 | Sigma-Aldrich | G-A-5 |
| IBA1 | 1:250 | Wako | PTP5154 |
| DAPI | 1:500 | Roche | - |
| In vivo | | | |
| Anti-mouse CD8α | 300µg, i.p. | BioXCell | 2.43 |
| Anti-mouse IgG2a Isotype | 300µg, i.p. | BioXCell | C1.18.4 |

**Table S2. RT-qPCR Primers**

| Target | Forward (5'-3') | Reverse (5'-3') | Species |
|---|---|---|---|
| GFAP | TCCTGGAACAGCAAAACAAG | CAGCCTCAGGTTGGTTTCAT | Mouse |
| IBA1 | GTCCTTGAAGCGAATGCTGG | CATTCTCAAGATGGCAGATC | Mouse |
| CD68 | ACTTCGGGCCATCTTTCTCT | GCTGGTAGGTTGATTGTCGT | Mouse |
| CD86 | ACGATGGACCCCAGATGCACCA | GCGTCTCCACGGAAACAGCA | Mouse |
| B-actin | CTGGCTCCTAGCACCATGAA | CGCAGCYCAGTAACAGTCCG | Mouse |

**Data S1. (separate file)**
Single Cell RNA Sequencing Data

**Data S2. (separate file)**
GO Ontology Analysis Data
